# Exploring Binary Relations for Ontology Extension and Improved Adaptation to Clinical Text

**DOI:** 10.1101/2020.12.04.411751

**Authors:** Luke T Slater, Robert Hoehndorf, Andreas Karwath, Georgios V Gkoutos

## Abstract

**Background:** The controlled domain vocabularies provided by ontologies make them an indispensable tool for text mining. Ontologies also include semantic features in the form of taxonomy and axioms, which make annotated entities in text corpora useful for semantic analysis. Extending those semantic features may improve performance for characterisation and analytic tasks. Ontology learning techniques have previously been explored for novel ontology construction from text, though most recent approaches have focused on literature, with applications in information retrieval or human interaction tasks. We hypothesise that extension of existing ontologies using information mined from clinical narrative text may help to adapt those ontologies such that they better characterise those texts, and lead to improved classification performance.

**Results:** We develop and present a framework for identifying new classes in text corpora, which can be integrated into existing ontology hierarchies. To do this, we employ the Stanford Open Information Extraction algorithm and integrate its implementation into the Komenti semantic text mining framework. To identify whether our approach leads to better characterisation of text, we present a case study, using the method to learn an adaptation to the Disease Ontology using text associated with a sample of 1,000 patient visits from the MIMIC-III critical care database. We use the adapted ontology to annotate and classify shared first diagnosis on patient visits with semantic similarity, revealing an improved performance over use of the base Disease Ontology on the set of visits the ontology was constructed from. Moreover, we show that the adapted ontology also improved performance for the same task over two additional unseen samples of 1,000 and 2,500 patient visits.

**Conclusions:** We report a promising new method for ontology learning and extension from text. We demonstrate that we can successfully use the method to adapt an existing ontology to a textual dataset, improving its ability to characterise the dataset, and leading to improved analytic performance, even on unseen portions of the dataset.

## Background

Biomedical ontologies provide sets of natural language labels and synonyms that constitute controlled domain vocabularies. These vocabularies are a fundamental resource for text mining [1]. However, ontologies also contain a wealth of semantic features, which enables their use in ontology-based analysis [2]. Recent approaches are increasingly bridging the gap between text mining, semantic analysis, and machine learning [3]. While many recent efforts have focused on the best ways to leverage the axioms that exist in ontologies to improve the power of analyses that use them, less attention has been given to whether text mining could be used to create additional classes and relationships in existing ontologies, particularly extending them in ways that could benefit analysis.

Ontology learning from text, however, has been an area of interest for many years [4, 5]. Most approaches focus on learning from literature, and their most common methods of validation are manual, or through use of consistency and satisfiability checks performed by automated reasoners [6]. Other evaluations include application integration, information retrieval, human interaction and performance against gold standard question-answer database [5]. However, the role of ontology learning for adaptation and extension of existing ontologies for improvement of classification tasks has not been previously explored, to our knowledge.

Previous approaches to ontology learning have used Stanford CoreNLP [7] for pre-processing and part-of-speech tagging [8, 9]. However, CoreNLP also includes an Open Information Extraction (OpenIE) algorithm, which can extract binary relations from text [10]. The relations it produces are derived from the syntax of sentences, often using the verb of the sentence as a predicate between two nouns or noun phrases which form a subject and object [11]. Some previous investigations have built OWL ontologies from literature text using these relations in combination with topic modelling approaches [12, 13]. However, these approaches were only applied to literature text, and have only been used to support applications of information retrieval and literature search. Furthermore, they focus on producing novel ontologies for a particular topic, rather than extending existing ontologies.

One study of using openIE for knowledge graph construction identified several open problems in the domain [14]:

### Entity linking

Linking named entities in the binary relation to identified entities in a knowledge graph.

### Entity Selection

Selecting the most optimal and relevant entity amongst candidates produced. For example, which substring of “aorta descent consistent with dissection” to use as the entity in the relation.

### Property linking

Similar to the problem of entity linking, linking the predicate in the binary relation to an ontology resource.

### Representation

How to model more expressive binary relations, extrapolating complex relations to several triples.

Instead of using the binary relationships extracted from text to construct an ontology, our approach uses an analysis of the components of extracted binary relationships to identify new classes that can be linked to existing ontology classes. This renders our approach relevant and applicable to the entity linking and entity selection problems. By adding new terms to an existing ontology in this fashion, we hypothesise that the ontology can be adapted to a text corpus, leading to its better characterisation through the annotation of more specific concepts. To explore this hypothesis, we developed a novel approach to ontology extension using triples produced by the OpenIE algorithm. We take a fully automated approach, with no manual curation or evaluation of correctness before final use, with the aim of producing an application ontology that is adapted to a particular dataset, and whose success can be measured by its role in improved classification performance. We apply the approach to a text corpus associated with MIMIC-III patient visits and generate an extended version of the Disease Ontology (DO) [15]. We describe the ontology, and then investigate its use for improving the performance of diagnosis extraction and classification from clinical text.

The triple mining and ontology construction modules are available as part of the Komenti semantic text mining framework, which is available under an open source licence at https://github.com/reality/komenti [16]. Files used for the construction of the MIMIC Adapted Disease Ontology (MADO), and for the automated validation are available at https://github.com/reality/mado_tripulate.

## Results

### Approach

We developed a method of extending existing ontologies with new classes mined from text. The resulting ontology facilitates the identification of more specific concepts expressed in the corpus text, in the context of the existing hierarchy. In order to extend an existing ontology, important entities in the text that are related to, but may extend, classes in the ontology of interest need to be identified. Building a vocabulary of relevant terms from the ontology and using it to identify sentences which contain mentions of ontology classes produces candidate sentences that may concern useful new terms.

This leaves the problem that we need to identify modified ontology terms that we can use to produce new classes. Binary relationship extraction algorithms identify relationships between entities in text. In this case, we consider the Open Information Extraction (OpenIE) algorithm, which uses syntactic analysis to identify entities and relationships. Because OpenIE works on this syntactic basis, and is therefore naive to the actual terms in the ontology, it is able to capture noun phrase entities which contain ontology terms, but extend them (make them more specific), allowing us to identify new classes that extend existing ontology classes. Furthermore, by examining only binary relations that contain at least one ontology term in both its subject and object, we can assume that these entities are both important to the text and relevant to the context of the ontology. The subject and object entity strings can then be decomposed into a combination of ontology terms and an optional specifying component. These new terms can then be added to the ontology, making them descendants of the classes they extend.

Once this has been done, the extended ontology can be used to annotate those more specific concepts in the text corpus. In this investigation, we do not make use of the binary relations themselves to add additional relationships to the ontology, beyond identifying classes to add to the hierarchy. The employment of binary relationship extraction in this case can be considered as a ‘filter,’ identifying important entities in the text to add to the ontology. Figure 1 describes the overall approach.

**Figure 1.**
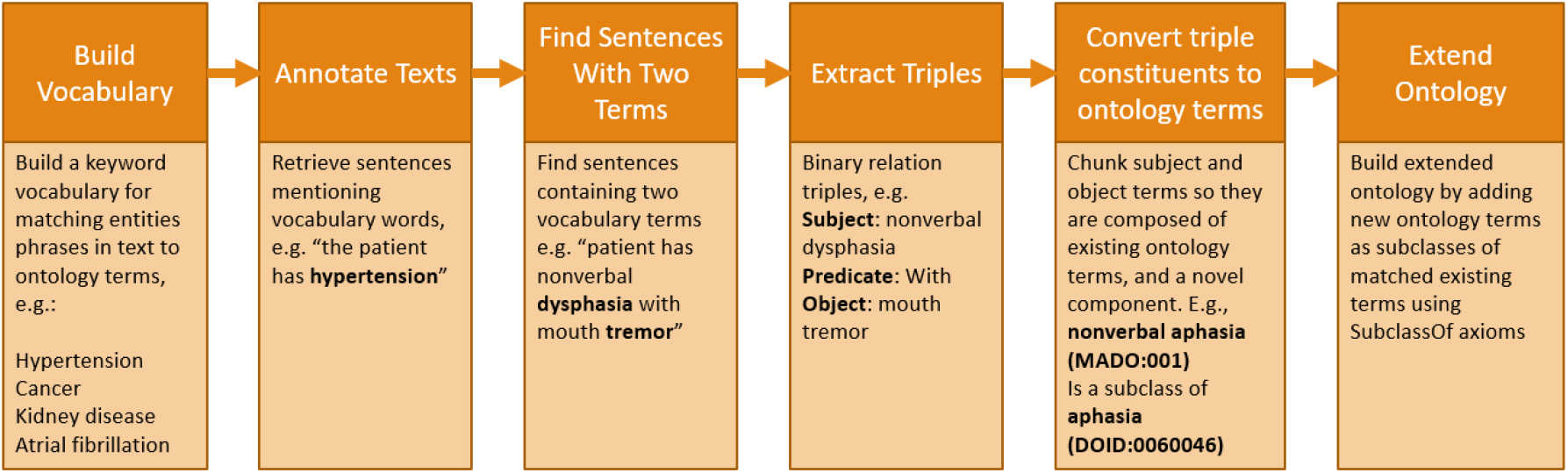
Description of our workflow to extract new classes from a corpus by annotating a text, and examining binary relations.

### Vocabulary Construction and Annotation

For the vocabulary construction and annotation stages, our approach uses Komenti’s existing pipeline for vocabulary construction and text annotation [16]. The vocabulary construction process uses the AberOWL reasoner-based ontology access framework [17] to build a list of labels and synonyms for a given input ontology. The annotation process uses this vocabulary to match mentions of the relevant concepts in text, using the Stanford RegexNER annotator implemented by CoreNLP. This outputs a list of annotated terms and the sentences that contain them. To identify sentences that could contain valid binary relations that pertain to terms in the input ontology, our algorithm then examines the list of annotations, filtering it to contain only sentences which contain at least two ontology terms.

### Entity Selection and Linking

The remaining sentences are then used as input to the Open Information Extraction (OpenIE) algorithm, which extracts binary relationships from input text. OpenIE produces triples, which represent a relationship between two entities in the form of a subject, predicate, and object. Since we are passing it sentences which contain two terms from our ontology, we intend to capture relationships which concern subjects and objects related to our ontology: either equivalent to a term, or containing terms that can be transformed into a new specifying class.

To integrate these new entities into our ontology, we need to describe them in terms of existing ontology classes. Subjects and objects are again annotated using the vocabulary that was previously used to identify candidate sentences. Subsequently a representation of the entities are produced by repeatedly identifying the longest annotation corresponding to an ontology term. When these have been exhausted, either the whole string has been consumed, or we are left with an arbitrary specifying string. In this way, we produce a new class, which we assert is a subclass of the existing ontology terms contained in the entity string. The algorithm performing this process is described in Figure 2.

**Figure 2.**
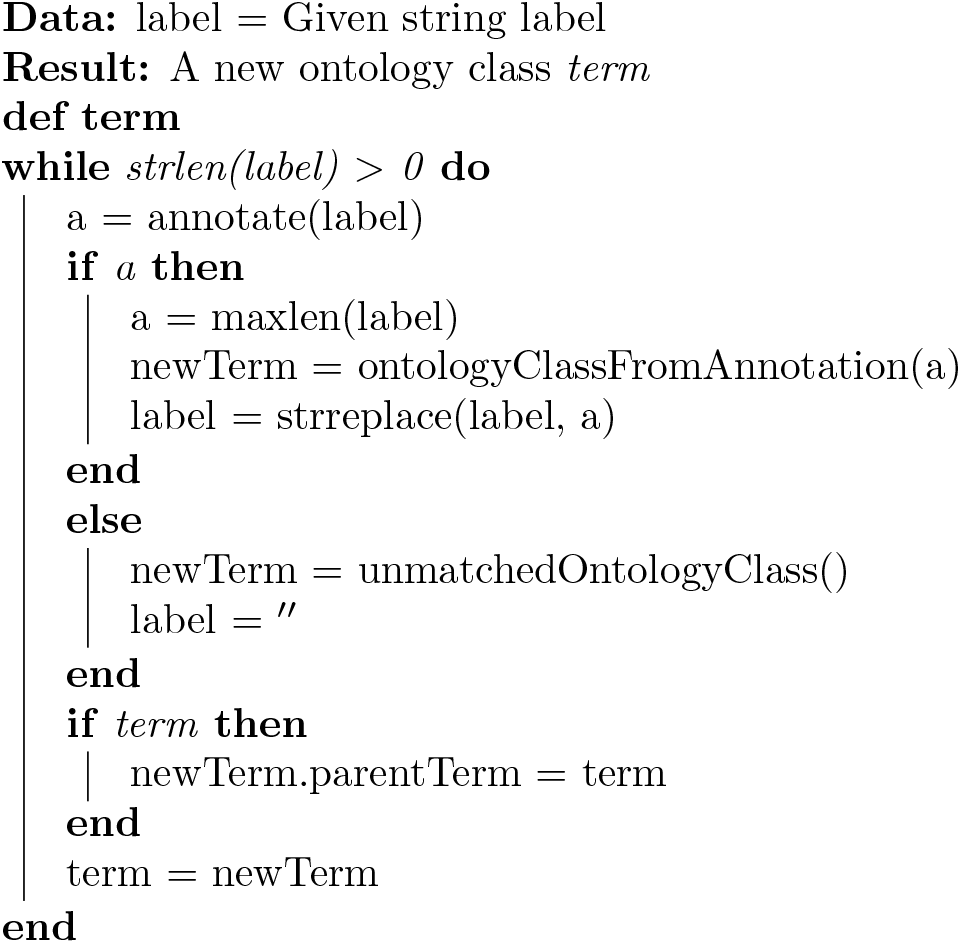
Algorithm for building hierarchical ontology terms from matched openIE labels by recursively consuming the label. The annotate function returns a list of ontology terms that were found in the input string.

For example, an entity label appearing as a subject or object in a triple could be “extreme hypertension.” Using a vocabulary constructed from the Disease Ontology (DO), the first annotation employed by the algorithm would match *hypertension* (DOID:10763). The algorithm would then recurse, attempting to annotate the remaining unconsumed string, ‘extreme.’ Since this word does not feature in DO, no annotation can be made. Thus, ‘extreme’ is noted as an unmatched concept, and we can produce a new class with the label ‘extreme hypertension,’ as a subclass of *hypertension* (DOID:10763). New terms can also be composed of multiple ontology classes, such as for the example ‘activation of apoptosis cell.’ Using a vocabulary constructed from the Gene Ontology, the algorithm would first identify *activation of apoptosis* (GO:0006915) as the annotation with the maximal string length. On the second round, it would match *cell* (GO:0005623), leading to a new ‘activation of apoptosis cell’ class, which would be a subclass of both *activation of apoptosis* and *cell*. This can lead to rich hierarchies, as other processed entities may specify further upon these intermediate classes, such as for ‘activation of apoptosis cell of mz-cha-1’ or ‘activation of apoptosis less than control.’ If no ontology term is matched at all, the whole string is returned as an unmatched concept, and is later thrown away by the algorithm, as it cannot be integrated with the ontology hierarchy.

### Ontology Extension

For each triple, after the subject and object have been processed, we add both to the list of new classes to add to the ontology, if we were able to construct an ontology term from both the subject and object. Otherwise, it is discarded. This is to ensure that we extend existing ontology structures and features; we would not be able to construct an ontology class from an entity label that does not contain at least one ontology term. Once all triples have been evaluated, a Komenti module is used to create a new ontology expressed by subclass axioms and labels, asserting the new classes in the context of the target ontology. Once this has been done, an import to the original ontology is added.

### MIMIC Adapted Disease Ontology

We applied our method to texts associated with 1,000 patient visits sampled from the MIMIC-III critical care database. Our input vocabulary contained terms from the Disease Ontology (DO) and the Phenotypic Trait Ontology (PATO) [18]. The DO terms were restricted to classes that contained cross references to ICD-9, which is the medical terminology used to annotate MIMIC-III patient visits in the structured data. We did this to reduce the running time of the experiment, by restricting our analysis to classes likely to be represented in the structured outcome data. We will refer to this as the ‘Trim DO’ vocabulary, to distinguish it from the ‘Full DO’ vocabulary, which contains all non-obsolete classes in the ontology. This produced the MIMIC Adapted Disease Ontology (MADO). The unmodified DO contains 18.591 classes, while MADO contains 20,818, meaning that the approach produced 2,227 new classes.

### Diagnosis Classification

To evaluate whether MADO could lead to an improved characterisation of the dataset, we compared its performance at classifying shared primary diagnosis over sets of MIMIC patient visits. We used the annotation terms as input to the Resnik semantic similarity algorithm, using the Best Match Average, and using a measure of information content derived from the probability of the annotation appearing in the entire set of annotations. We then evaluated our approach’s performance at identifying a shared primary diagnosis when ranking patient visits by their similarity scores.

Our evaluation included several settings, spanning different combinations of vocabularies used as input, and ontologies used to calculate the similarity scores. Since a Trim DO vocabulary (the subset with ICD-9 cross mappings) was used to produce MADO, we also included results for use of the Trim DO vocabulary without extensions to ensure that improved performance did not derive from use of the slimmer vocabulary alone.

**Table 1.** Performance for matching first diagnosis of MIMIC patients with different combinations of annotation vocabularies and ontologies used for semantic similarity score calculation.

| Setting | Vocabulary | Ontology | MRR | MAP | Precision | p-value (<) |
| --- | --- | --- | --- | --- | --- | --- |
| Source-1000 | Full DO | DO | 0.939 | 0.844 | 0.937 | - |
|  | Trim DO | DO | 0.947 | 0.88 | 0.917 | 0.01466 |
|  | MADO | MADO | **0.995** | **0.916** | **0.992** | 6.458e-05 |
| Unseen-1000 | Full DO | DO | 0.931 | 0.841 | 0.929 | - |
|  | Full DO | MADO | 0.88 | 0.795 | 0.876 | 0.007279 |
|  | Trim DO | DO | 0.958 | 0.885 | 0.932 | 0.0001161 |
|  | Trim DO | MADO | 0.958 | 0.887 | 0.933 | 0.003315 |
|  | MADO | MADO | **0.986** | **0.92** | **0.981** | 0.009601 |
| Unseen-2500 | Full DO | DO | 0.943 | 0.842 | 0.941 | - |
|  | Full DO | MADO | 0.896 | 0.798 | 0.891 | 2.2e-16 |
|  | Trim DO | DO | 0.932 | 0.865 | 0.898 | 0.004658 |
|  | Trim DO | MADO | 0.933 | 0.866 | 0.899 | 8.035e-05 |
|  | MADO | MADO | **0.986** | **0.913** | **0.98** | 2.2e-16 |

## Discussion

Our results show that the developed method was successful in constructing a MIMIC adapted disease ontology. The results of the characterisation evaluation demonstrate that the use of the MADO ontology and vocabulary consistently out-performs two different configurations of DO vocabulary used with DO and MADO for classificaiton of primary diagnosis. Furthermore, not only does our method improve characterisation based on text the ontology that was learned upon, it also improves characterisation from unseen text in the same corpus. This provides evidence that this process of training and extending an ontology on a dataset adapts that ontology to it, increasing its ability to characterise the dataset.

One limitation of the study is that it only considers primary diagnosis. Our evaluation could be considered a simplified version of those presented in several other investigations which use various learning methods to assign ICD-9 codes based on MIMIC-III text [19]. Our work only assigns the primary diagnosis, as a method of evaluating improvement between two settings. To make the work more practical, and to further evaluate the approach, we could consider transforming the task to classify several codes. However, this presents several challenges, primarily due to semantic similarity being an unsupervised and unguided approach. However, we believe that the similarity approach should capture this information, e.g. that patients with a secondary diagnosis of ‘pain’ would be more similar to other patients with a pain diagnosis. The challenge is in developing a decision procedure for making that classification, while overlooking other reasons (i.e. a different shared diagnosis) that patients may be highly similar. The advantages of developing this kind of decision procedure would be the ability to use this method for a more complete characterisation of the dataset, as well as the ability to directly compare performance with other approaches using the same evaluation methods. A follow-up investigation could measure how well annotations predicted overall patient diagnosis structure. These methods could also be applied to other benchmark issues that have typically been solved with similarity-based approaches, such as gene enrichment or protein-protein similarity. A future validation could also work to determine whether larger training sets could lead to a greater validation performance, and whether this value could be established as a hyper-parameter set based on size or other information concerning a target corpus.

Nevertheless, the evaluation showed that our approach improved the performance of the task of classifying primary diagnosis from text. This has potential applications in the clinical space, including automated diagnosis and classification of patients from text profiles. Previous work involved the use of semantic similarity over the Human Phenotype Ontology for differential diagnosis of medical conditions [20–22]. However, these phenotypic profiles are usually produced by experts, or by patients themselves. We would like to investigate whether it is possible to construct these profiles from text, particularly aided by methods that can construct and adapt multiple ontologies around particular diseases, datasets, and relevant entities.

While our work shows that the approach can adapt an individual ontology to a text corpus for improved classification, this work could potentially have other uses and implications. As well as extending other single ontologies to better characterise other text corpora with application to other classification problems, the approach also has the potential to capture relationships between different domain ontologies. While this investigation only involves DO, PATO, and the classes they include, the algorithm would also work to connect more ontologies that describe different domains. For example, a vocabulary with terms from DO and GO would be capable of producing new classes that combined genetic components and diseases, such as in the case of ‘cancer cell.’ This would potentially allow for the integration of information from different domains contributing to singular tasks. In the case of semantic similarity, involving several ontologies produces an additional level of complexity [23].

Furthermore, while the approach employs the use of binary relation extraction, it does not currently make use of the relationships themselves. Observed relationships from text could be used as ABox axioms, defining observations of instances of ontology classes in text. Semantic similarity approaches do not make use of such information, although recent approaches to machine learning with ontologies, such as OPA2Vec [24] could take these relationships into account. This would potentially further increase the power of cross-ontology and integrated domain analysis, as relationships could be established between classes of different ontologies through observations as well as class composition.

Both the construction of the ontology, and thereby analysis performance could be affected by investigating the use of other Komenti features, particularly those for negation detection and synonym expansion [25, 26]. In the former case, the evaluation of negation could prevent incorrect or explicitly negated facts from being used in the produced knowledgebase, while the latter case has been shown to improve overall characterisation of text.

## Conclusions

Our work describes a new approach to extending existing ontologies with new classes through analysis of text corpora. The ontology extensions essentially adapt the ontology to better suit the text corpus, and leads to improved classification performance using the extended set of annotations, even on unseen portions of the dataset. We believe that the approach is promising, and could have further implications, which we intend to explore in future work.

## Methods

The algorithm was developed using the Komenti semantic text mining framework, which is available under an open source licence at http://github.com/reality/Komenti [16]. It makes use of Stanford CoreNLP [7], and particularly the RegexNER annotator for identifying entities in text, and the openIE algorithm for extracting triples from text.

### Corpus Generation

Texts were sourced from the MIMIC-III critical care database [27]. MIMIC is a publicly available dataset describing over 50,000 critical care visits, including natural language text linked with many structured features. We sampled three sets of patient visits from MIMIC: one of 1,000 patient visits for learning the ontology, and test sets of 1,000 and 2,500 patient visits. Patient visits were associated with their primary diagnosis, the first listed for the record in the ICD9 DIAGNOSES table. These codes are produced by clinical coding specialists, by examining the texts associated with the visit. We then concatenated all text records for each patient visit from the NOTEEVENTS table into one text file, and pre-processed the text to remove newlines, improve sentence delineation, and lemmatise words.

We used two base vocabularies for our experiments. The first, referred to as ‘Full DO’ throughout the article, contained all labels and synonyms in the Disease Ontology (DO) [15]. The second was limited to labels and synonyms in 2,118 DO terms that were annotated with a database cross-reference to ICD-9 used in an effort to reduce noise from terms not represented in ICD-9. We obtained the unexpanded and expanded synonyms for these terms on 08/07/2020. Both sets of labels were also lemmatised.

### Ontology Extension

Our ontology extension approach was applied to the pre-selected training set of 1,000 patient visits, using the Trim DO vocabulary. The produced ontology was then merged with DO, by copying all axioms from DO into the new ontology. This merged ontology is referred to as the MIMIC Adapted Disease Ontology (MADO). The ontology constructor also contains an optional flag that outputs a new Komenti-compatible vocabulary with the novel classes and pre-existing ontology classes used as components in the ontology extension. This feature was used to create a new vocabulary, which was concatenated with the Trim DO vocabulary, and used in the experiments that refer to the MADO vocabulary.

### Evaluation

Komenti was used to annotate the text associated with each patient visit. We then used the set of terms associated with it to produce a semantic similarity matrix for patient visits, using the Resnik measure of pairwise similarity for each annotated term [28], normalised into a groupwise measure using the best match average method [29]. Information content was calculated using the probability of the term appearing as an annotation in the totality of the set of annotations [28]. The similarity matrix was computed using the Semantic Measures Library [30].

We evaluated the similarity matrix using mean reciprocal rank, mean average precision, and precision to measure performance in predicting shared primary patient diagnosis. A true case was considered to be whether a pair of patient visits had the same primary diagnosis (as per the MIMIC-III database). For mean average precision, we considered only the 10 most similar patients for each patient, establishing a cut-off to limit the effect of patients who were ranked more similar for reasons of other diagnosis. The p-value was calculated using the built-in *wilcox*.*test* function of R version 3.4.4 [31].

## Author’s contributions

LTS conceived of the study, performed the experiments, implemented the software, and wrote the first draft of the manuscript. LTS, RH, and AK contributed to the experimental design. RH and GVG contributed to the manuscript. GVG and RH supervised the project. All authors contributed to the interpretation of the results, as well as revised and approved the manuscript for submission.

## Acknowledgements

We would like to thank Dr Egon Willighagen, and Dr Paul Schofield for helpful discussions surrounding described investigations.

